# Deep learning-based quantification of arbuscular mycorrhizal fungi in plant roots

**DOI:** 10.1101/2021.03.05.434067

**Authors:** Edouard Evangelisti, Carl Turner, Alice McDowell, Liron Shenhav, Temur Yunusov, Aleksandr Gavrin, Emily K. Servante, Clément Quan, Sebastian Schornack

## Abstract

Soil fungi establish mutualistic interactions with the roots of most vascular land plants. Arbuscular mycorrhizal (AM) fungi are among the most extensively characterised mycobionts to date. Current approaches to quantifying the extent of root colonisation and the abundance of hyphal structures in mutant roots rely on staining and human scoring involving simple, yet repetitive tasks prone to variations between experimenters.

We developed AMFinder which allows for automatic computer vision-based identification and quantification of AM fungal colonisation and intraradical hyphal structures on ink-stained root images using convolutional neural networks.

AMFinder delivered high-confidence predictions on image datasets of roots of multiple plant hosts (*Nicotiana benthamiana*, *Medicago truncatula*, *Lotus japonicus*, *Oryza sativa*) and captured the altered colonisation in *ram1-1*, *str* and *smax1* mutants. A streamlined protocol for sample preparation and imaging allowed us to quantify mycobionts from the genera *Rhizophagus*, *Claroideoglomus*, *Rhizoglomus* and *Funneliformis* via flatbed scanning or digital microscopy including dynamic increases in colonisation in whole root systems over time.

AMFinder adapts to a wide array of experimental conditions. It enables accurate, reproducible analyses of plant root systems and will support better documentation of AM fungal colonisation analyses. AMFinder can be accessed here: https://github.com/SchornacklabSLCU/amfinder.git

## Introduction

Soil fungi establish mutualistic interactions with the roots of more than 85% of vascular land plants (Brundrett & Tedersoo, 2018). These interactions, termed mycorrhizae, lead either to the formation of a dense hyphal sheath surrounding the root surface (ectomycorrhizae) or to fungal hyphae penetrating host tissues (endomycorrhizae) (Brundrett, 2007). The best-characterized type of endomycorrhiza, called arbuscular mycorrhiza (AM), involves species from the subphylum Glomeromycotina (Schüßler *et al.*, 2001; Spatafora *et al.*, 2016). AM fungal hyphae grow toward plant roots following the exchange of diffusible chemical cues (Luginbuehl & Oldroyd, 2017). At root surface penetration points, hyphae differentiate into swollen or branched structures termed hyphopodia. Following entry and crossing of the root epidermis, hyphae spread between cortical cells (*Arum*-type colonization) or via intracellular passages of cortical cells (*Paris*-type colonization) (Dickson, 2004). The differentiation of highly-branched intracellular exchange structures, the arbuscules, accompanies hyphal growth and enables a reciprocal transfer of nutrients between symbionts (Luginbuehl & Oldroyd, 2017). Post-arbuscular development includes the differentiation of vesicles and spores. While these successive differentiation events reflect a precise morphogenetic program, the whole hyphal network is not synchronized. As a result, the various types of intraradical hyphal structures occur simultaneously inside plant roots (Montero *et al.*, 2019).

*Rhizophagus irregularis* (formerly *Glomus intraradices*) is one of the most extensively characterised mycobionts in endomycorrhiza research. To date, genetic manipulation of *R. irregularis* remains challenging (Helber & Requena, 2008) and the main advances in AM fungal symbiosis research relate to the experimentally more tractable plant hosts. The extent of root colonisation and the relative abundance of intraradical hyphal structures in mutant roots are essential parameters for characterising host genes that underlie mycorrhiza establishment and accommodation (Montero *et al.*, 2019). Mycorrhiza-responsive host genes facilitate the molecular quantification of fungal colonisation. For instance, expression of the *Medicago truncatula Phosphate transporter 4* (*MtPT4*) gene is limited to the root tip in the absence of mycorrhiza (Volpe *et al.*, 2016), while cells with arbuscules express *MtPT4* to enable plant acquisition of inorganic phosphate (Harrison *et al.*, 2002; Maeda *et al.*, 2006; Javot *et al.*, 2007). Likewise, the abundance of transcripts encoding *M. truncatula* Blue Copper-Binding Protein 1 and *Lotus japonicus* apoplastic subtilase SbtM correlate with stage transitions during arbuscule development (Hohnjec *et al.*, 2005; Takeda *et al.*, 2009; Parádi *et al.*, 2010). Complementary to molecular methods and independent of gene sequence knowledge is the visual diagnosis of AM fungal colonisation. It involves differential staining of fungal cell walls (Vierheilig *et al.*, 1998, 2005; Hulse, 2018) followed by random sampling and counting using a grid-intersect method (Giovannetti & Mosse, 1980). This method is considered a standard in mycorrhiza research (Sun & Tang, 2012).

Deep learning encompasses an extensive class of computational models that learn to extract information from raw data at multiple levels of abstraction, thereby mimicking how the human brain perceives and understands information (Voulodimos *et al.*, 2018). In supervised learning problems, where example data labelled with correct outputs are available, these models can be iteratively improved to minimise discrepancies between correct and model-predicted outputs considering all possible interfering factors (O’Mahony *et al.*, 2020). With the increase in computing power over recent years, deep learning has fostered tremendous data analysis advances. Computer vision is one of the most iconic examples, with the development of convolutional neural networks (CNNs), a class of deep learning methods inspired by models of the visual system’s structure (LeCun *et al.*, 1998). A basic image classification CNN begins with a stack of convolutional and pooling layers, each providing the input to the next; these allow detection of increasingly large and complex features in the input image whilst preserving the grid-like nature of the input (Voulodimos *et al.*, 2018; Dhillon & Verma, 2020). This is followed by one or more fully connected layers of neurons which take the resulting feature map and implement the high-level reasoning needed to classify the image. CNNs underlie breakthrough advances in diverse technological and biomedical domains including face recognition, object detection, diagnostic imaging, and self-driving cars (Matsugu *et al.*, 2003; Szarvas *et al.*, 2005; Bojarski *et al.*, 2016; Yamashita *et al.*, 2018). CNNs also benefit botany by enabling the identification of flowers (Liu *et al.*, 2017) and ornamental plants (Sun *et al.*, 2017), while CNN-based plant pathology diagnostic tools identify crop diseases based on leaf symptoms (Mohanty *et al.*, 2016; Ferentinos, 2018; Thapa *et al.*, 2020).

We took advantage of CNNs to develop the Automatic Mycorrhiza Finder (AMFinder), an automatic, user-supervised tool suite for *in silico* analysis of AM fungal colonisation and recognition of intraradical hyphal structures. Using AMFinder, we quantified fungal colonisation dynamics on whole *Nicotiana benthamiana* root systems using low-resolution, flatbed scanner-acquired images of ink-stained roots. AMFinder accurately quantified changes in the extent of *R. irregularis* colonisation in *M. truncatula ram1-1*, *str*, and *O. sativa smax1* plant mutants compared to their wild type lines. Moreover, AMFinder robustly identified colonised root sections and intraradical hyphal structures in several plant species commonly used in mycorrhiza research, including *M. truncatula*, *L. japonicus*, and *O. sativa*, and is compatible with the AM fungi *Claroideoglomus claroideum*, *Rhizoglomus microaggregatum*, *Funneliformis geosporum* and *Funneliformis mosseae*. We developed command-line prediction tool paired with a standalone graphical browser to enable efficient browsing of large images and manual curation of computer predictions. Overall, our work provides a framework for reproducible automated phenotyping of AM fungal colonisation of plant roots.

## Materials and methods

### Plant material

*Nicotiana benthamiana* is a laboratory cultivar obtained from The Sainsbury Laboratory, Norwich, UK, originating from Australia (Bally *et al.*, 2018). *Medicago truncatula* R108 seeds were provided by Giles Oldroyd (The Sainsbury Laboratory, UK). *Lotus japonicus* Gifu seeds were provided by Simona Radutoiu (Aarhus University, Denmark). Rice (*Oryza sativa* subsp. *japonica*) plant material was described elsewhere (Choi *et al.*, 2020).

### Seed germination

*N. benthamiana* seeds were germinated on Levington F2 compost (ICL, Ipswich, UK) for one week at 24°C with a 16-h photoperiod. *M. truncatula* seeds were scarified in sulphuric acid for 5 min, rinsed in sterile water and surface-sterilized in bleach for 5 min. Seeds were then soaked in water for 30 min and stratified for 3 days at 4°C in the dark. *L. japonicus* seeds were scarified with sandpaper, surface-sterilized in bleach for 15 min and soaked overnight in water at 4°C. Germination was induced at 20°C. *O. sativa* seed germination was described elsewhere (Choi *et al.*, 2020).

### Growth conditions for AM colonisation

One-week-old seedlings were transferred to 6×5 cellular trays containing silver sand supplemented with a 1:10 volume of AM fungal crude inoculum and grown at 24°C with a 16-h photoperiod. *R. irregularis*, *C. claroideum* and *F. geosporum* crude inocula were obtained from PlantWorks (Sittingbourne, UK). *F. mosseae* crude inoculum was obtained from MycAgro (Bretenière, France). *N. benthamiana* plants were watered with a low-phosphate Long Ashton nutrient solution (Hewitt, 1966), while milliQ water was used for *L. japonicus* and *M. truncatula* plants. *O. sativa* growth conditions and AM colonisation conditions were described elsewhere (Choi *et al.*, 2020). Plant roots were harvested at either 4 or 6 weeks post-inoculation and directly used for staining or total mRNA extraction.

### Fungal staining

A modified ink-vinegar method (Vierheilig *et al.*, 1998) was used to stain fungal structures within plant roots. Briefly, roots from 4- and 6-week-old plants were incubated in 10% (w/v) potassium hydroxide (KOH) for 10 min at 95°C and rinsed in 5% (v/v) acetic acid before staining in staining solution (5% (v/v) Sheaffer Skrip black ink, 5% (v/v) acetic acid) (A.T. Cross Company, Providence, RI, USA) for 10 min at 95°C. Stained roots were rinsed in distilled water, followed by clearing in ClearSee (Kurihara *et al.*, 2015) for 1 min **(Fig. S1)**. Cleared roots were mounted in a glycerol-containing mounting medium (20% (v/v) glycerol, 50 mM Tris-HCl pH 7.5, 0.1% (v/v) Tween-20).

### Scanning and bright field imaging

Low-magnification images of ink-stained roots were acquired with an Epson Perfection flatbed scanner (Epson UK, Hemel Hempstead, UK) using default settings and a resolution of 3200 dots per inch **(Fig. S2)**. High-magnification images were acquired with a VHX-5000 digital microscope (Keyence, Milton Keynes, UK) equipped with a 50-200× zoom lens set to 200× magnification, using transillumination mode and focus stacking. Images of *M. truncatula* arbuscules were obtained with an Axio Imager M2 (Zeiss, Germany) microscope equipped with a 64× NA 1.4 oil immersion objective lens using differential interference contrast (DIC) illumination.

### Quantitative RT-PCR

Total RNA was extracted from 100 mg root material using RNEasy Mini Kit (Qiagen, Hilden, Germany) according to the manufacturer’s instructions. Quality was assessed by electrophoresis on an agarose gel. One microgram was reverse transcribed to generate first-strand complementary DNA (cDNA) using IScript cDNA Synthesis kit (Bio-Rad, Hercules, CA, USA). qRT-PCR experiments were performed with 2.5 μl of a 1:20 dilution of first-strand cDNA and LightCycler 480 SYBR Green I Master mix, according to the manufacturer’s instructions (Roche, Basel, Switzerland). Gene-specific oligonucleotides were designed with BatchPrimer3 software (You *et al.*, 2008), and their specificity was validated by analysing dissociation curves after each run. Genes encoding L23 (Niben101Scf01444g02009) and FBOX (Niben101Scf04495g02005) were selected as constitutive internal controls for *N. benthamiana* (Liu *et al.*, 2012). Primers for transcripts encoding RiEF1a and NbBCP1b (Niben101Scf07438g04015.1) are listed in **Table S1**. Six biological replicates of the entire experiment were performed. Gene expression was normalised with respect to constitutively expressed internal controls (Vandesompele *et al.*, 2002) and plotted using R (https://www.r-project.org/).

### Software design

#### Implementation

AMFinder implements a semi-automatic pipeline for quantification of fungal colonisation and intraradical hyphal structures in root pictures. It comprises a command-line program (`amf`) for root image analysis and a standalone interface (`amfbrowser`) for user supervision and validation of `amf` predictions **(Fig. 1)**. `amf` is written in Python (https://www.python.org/) (van Rossum & Drake, 2009; Srinath, 2017) and uses the widely used Tensorflow (https://www.tensorflow.org/) and Keras (https://keras.io/) machine learning libraries (Chollet & others, 2015; Abadi *et al.*, 2016). `amfbrowser` is written in OCaml (https://ocaml.org/) (Leroy *et al.*, 2020) using the 2D graphics library Cairo (https://www.cairographics.org/) and the cross-platform widget toolkit GTK (https://www.gtk.org/). Both programs communicate through a standard ZIP archive file that stores amf probabilities, user annotations, and image settings. AMFinder deploys on Microsoft Windows, macOS and GNU/Linux.

**Figure 1.**
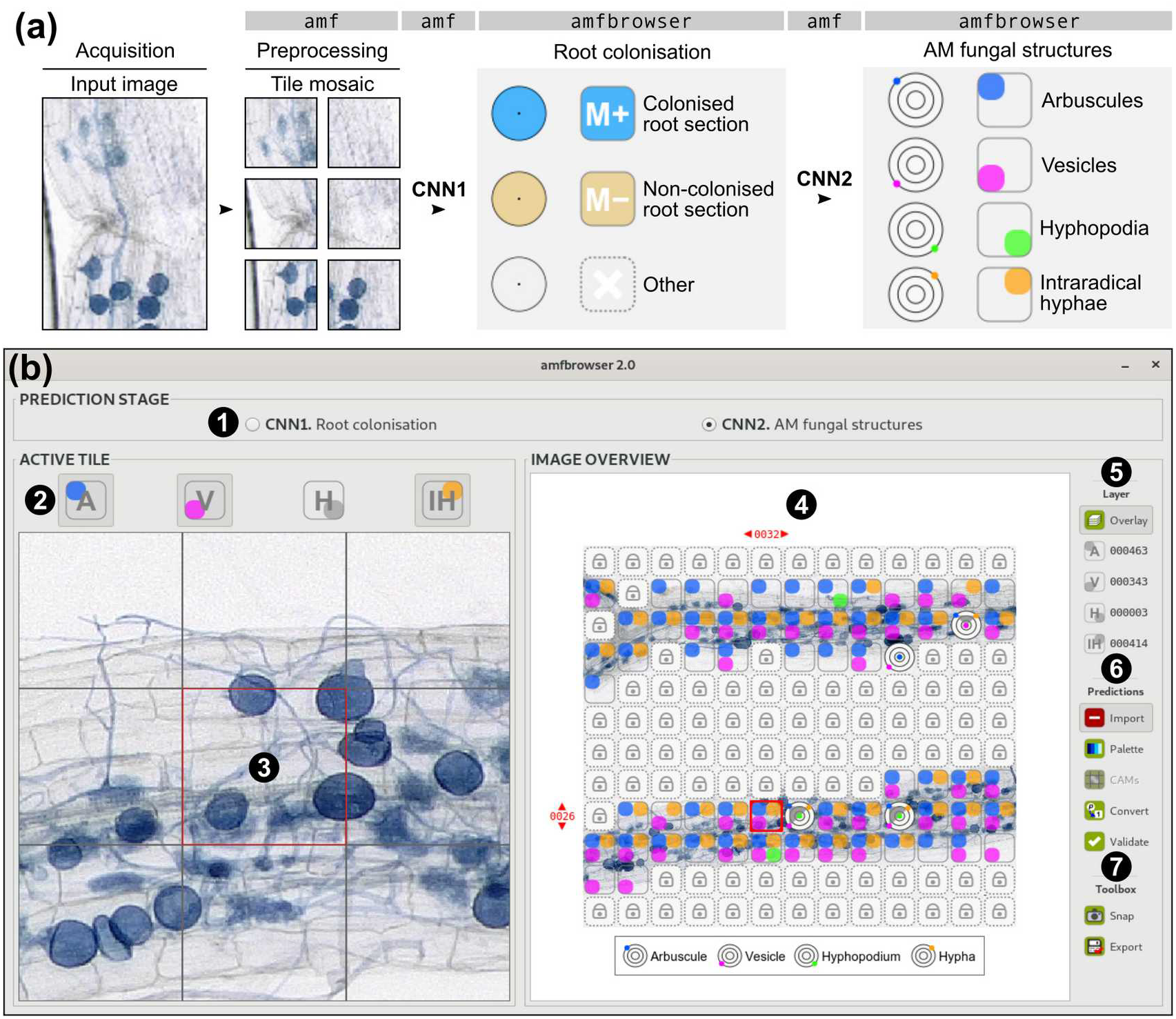
AMFinder enables a semi-automated, user-supervised analysis of AM fungal colonisation *in silico*. **(a)** AMFinder uses a two-stage prediction pipeline for image annotation. First, input images are split into tiles and processed by `amf` neural network 1 (CNN1) to identify colonised root sections. Computer predictions are displayed as pie charts and are converted to annotations under user supervision. If resolution allows, tiles corresponding to colonised areas can be further analysed by `amf` CNN2 to identify intraradical hyphal structures. The probabilities of occurrence of the different structures are shown in a radar plot. **(b)** Representative screenshot of an `amfbrowser` annotation session. ❶ Buttons to switch the display between prediction stages. ❷ Clickable buttons to define the annotations present in the active tile. ❸ Magnified view of the active tile (red square) and eight surrounding tiles. ❹ Annotation mosaic overview. ❺ Layer toolbar to filter the display. Numbers indicate annotation counts for the whole image. ❻ Prediction toolbar to load predictions, fix ambiguous cases and generate annotations. ❼ Export functions.

#### Analysis pipeline

The AMFinder pipeline consists of three or five steps depending on image resolution. First, image segmentation is performed by `amf` during an initial pre-processing step. Images are divided into square tiles using a user-defined tile size depending on image magnification and resolution **(Fig. 1a)**. Next, tiles are analysed individually to label colonised root segments **(Fig. 1a)**. The third step of the analysis consists of the user-supervised conversion of `amf` predictions (i.e. probabilities) to annotations using `amfbrowser` **(Fig. 1a, b)**. If resolution allows, a second round of `amf` predictions can be achieved to assess the occurrence of intraradical hyphal structures on colonised tiles only. As for the first round of predictions, computer-generated probabilities are then converted to annotations under user supervision using `amfbrowser` **(Fig. 1a, b)**.

### Deep learning

#### Classifier design

Identifying AM fungal colonisation and intraradical hyphal structures on images of stained roots constitute a computer vision problem that is efficiently solved by convolutional neural networks (CNNs). AMFinder implements two independent CNN-based classifiers to predict colonisation (CNN1) and intraradical hyphal structures (CNN2) **(Fig. S3)**. As the features we are interested in are mainly small, local structures, CNN1 was chosen to be a fairly shallow network. It comprises four blocks of 3×3 filters (convolutions) interleaved with size-reduction layers (maximum pooling) **(Fig. S3a)**, followed by a classifier made of three fully connected layers which compute the probabilities of each tile belonging to the mutually exclusive classes ‘colonised root section’ (M+), ‘non-colonised root section’ (M−), and ‘background/not a root/other’ **(Fig. 1a, Fig. S3b)**.

The CNN2 classifier predicts the occurrence of arbuscules (A), vesicles (V), hyphopodia (H), and intraradical hyphae (IH) on tiles labeled as ‘colonised root section’ (M+) during CNN1 analysis. Its architecture is essentially the same as that of CNN1 **(Fig. S3a)**. However, the probability that each different type of intraradical hyphal structure is present is computed by a separate stack of three fully connected layers atop the convolutional and pooling layers **(Fig. S3c)**. This design was chosen to facilitate future extensions of AMFinder, for instance to predict other types of structures (e.g. spores) or variations of a given structure (e.g. different degrees of arbuscule branching), by allowing additional expressive power to be quickly added to the network.

#### Datasets

The CNN1 training dataset comprised 32 images of ink-stained, ClearSee-treated *N. benthamiana* roots colonised with *R. irregularis*, acquired using either a flatbed scanner (5 images) or a digital microscope (27 images), yielding 175105 tiles annotated as follows: 15364 tiles belonged to the ‘colonised root section’ annotation class, 19455 tiles corresponded to non-colonised roots, and 140286 tiles consisted of background, small debris and air bubbles **(Table S2)**.

The CNN2 training dataset consisted of 55 high-resolution images of colonised *N. benthamiana* roots prepared as above. It comprised 20564, 16420, and 25077 tiles containing arbuscules, vesicles and intraradical hyphae, respectively. Only 508 tiles contained hyphopodia, preventing us from achieving efficient hyphopodia training due to the scarcity of this hyphal structure **(Table S3)**.

A bootstrap method (a highly simplified type of uncertainty sampling as used in active learning) was used to annotate images. First, ten images were manually annotated through `amfbrowser` and used to train a development (pre-alpha) version of AMFinder CNNs. Trained CNNs were then used to annotate the remaining images. Computer predictions were individually inspected through `amfbrowser` to correct certain low-confidence predictions (namely those with all network outputs below 0.5).

Datasets were randomly split into two subsets for training and validation in an 85:15 ratio. When data augmentation was used, the tiles in the training set were augmented by randomly applying the following modifications: horizontal flips, vertical flips, brightness adjustment (± 25%), colour inversion (image negation), desaturation (to create a grayscale image), or random adjustment of hue (rotating colours around a colour wheel).

An independent test set was used for independent assessment of CNN capabilities **(Table S4)**. It consisted of 20 manually-annotated images of 81 tiles each, acquired with either a flatbed scanner (10 images, used with CNN1 only) or a digital microscope (10 images, used for both CNNs).

#### Classifiers parameters and training

Both classifiers were trained for 100 epochs (i.e. complete training cycles) with a batch size of 32 during both the initial bootstrapping stage described above and the main training stage. For optimal training, background over-representation was compensated by randomly removing excess background tiles. Training weights were assigned based on tile count in each annotation class to account for any residual imbalance. Consistent with their respective output, categorical cross-entropy was used as the CNN1 loss function, while binary cross-entropy was used for each CNN2 classifier (Gordon-Rodriguez *et al.*, 2020). To prevent overfitting, an early stopping mechanism was used to prematurely terminate training and to restore the best-performing model parameters if the loss did not decrease for twelve training cycles in a row. Both classifiers relied on the Adam optimizer (Kingma & Ba, 2015) with an initial learning rate of 0.001 which was exponentially decayed with a factor of 0.9. In addition, learning rate was further decreased to one fifth if the loss did not decrease for two successive epochs, until it reached a minimum rate of 10^−6^. AMFinder training was achieved on a High-Performance Computing (HPC) system running Linux Ubuntu, using 10 cores and 20 Gb RAM.

#### Classifiers evaluation

Three parameters were determined to evaluate the classifier results. The accuracy is given by the relation:

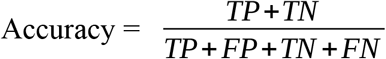

where TP (true positive rate) indicates accurate positive identifications (e.g. a tile containing intraradical hyphal structure is identified as colonised), TN (true negative rate) indicates accurate negative identifications (e.g. a tile containing root tissues only is identified correctly as non-colonised), FP (false positive rate) indicates that the observation is different but predicted as true (e.g. a non-colonised root tile is identified as colonised), and FN (false negative rate) indicates that a true observation is predicted to be different (e.g. a colonised root tile is identified as non-colonised). Accuracy counts all kinds of errors with the same costs and classifiers were thus further analysed using sensitivity and specificity:

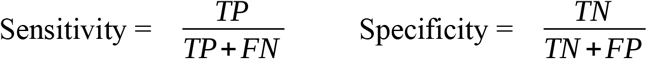

A high sensitivity indicates that the annotation class is correctly recognised (i.e. there are few false negatives), while high specificity indicates that a high number of tiles not labelled with a given annotation class *x* indeed do not belong to class *x* (i.e. there are few false positives).

## Results

### AMFinder robustly identifies intraradical structures in AM fungi

Convolutional neural network (CNN)-based classifiers require training to recognise the desired image classes. We trained the AMFinder CNNs using images of ink-stained roots of *N. benthamiana* plants inoculated with *R. irregularis* (Fig. 2a, b, Tables 1, 2). To improve CNN generalisation and reduce overfitting, we independently trained CNNs with augmented data **(Fig. 2c, d, Tables 1, 2)**. We then assessed CNNs performance on an independent test dataset **(Tables 1, 2, S4)**. CNN1 labelled colonised (M+) and non-colonised (M−) root sections, and background (Other) with an overall accuracy of 97% **(Fig. 2a, b, Table 1)**. The classifier performed similarly on low- **(Fig. 2a)** and high-magnification **(Fig. 2b)** images. Training on augmented data (CNN1v2) reduced the overall performance of the network to 94% when evaluated on ink-stained images **(Table 1)**. However, CNN1v2 was able to accurately predict colonisation on grayscale and inverted images **(Fig 2.d)**, suggesting network generalisation was improved compared to CNN1v1.

**Figure 2.**
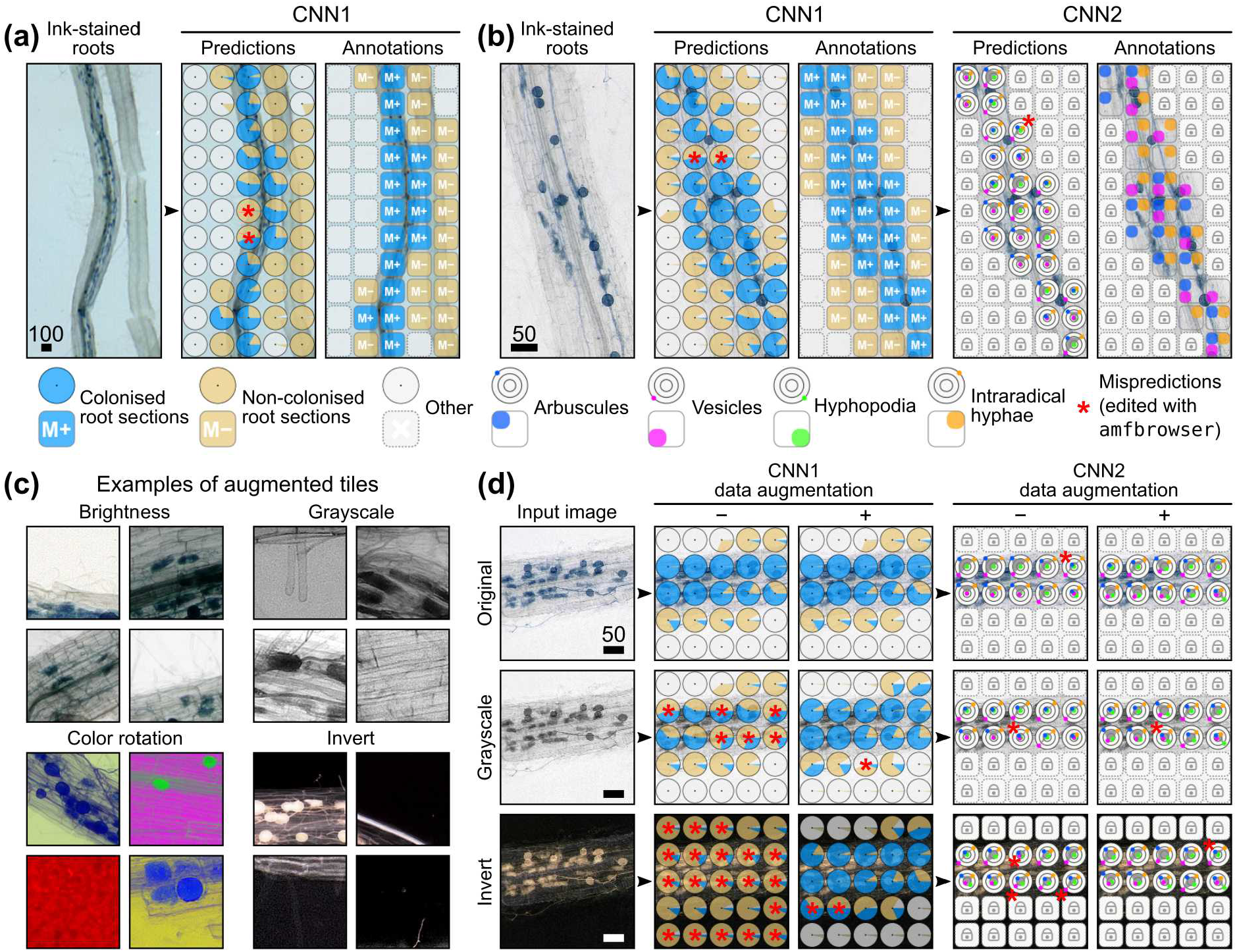
AMFinder accurately labels *Rhizophagus irregularis* colonisation and intraradical hyphal structures in *Nicotiana benthamiana* roots. **(a)** Schematic view of CNN1 predictions on a low-resolution image of ink-stained *N. benthamiana* roots acquired with a flatbed scanner. **(b)** Schematic view of CNN1 (left) and CNN2 (right) predictions on a high-resolution image of ink-stained *N. benthamiana* roots acquired with a digital microscope. **(c)** Examples of augmented tiles used for CNN1 and CNN2 training. **(d)** Comparison of CNN1 and CNN2 performance on images of ink-stained *N. benthamiana* roots colonised with *R. irregularis*, either unaltered, desaturated (grayscale), or inverted. Mispredictions (red asterisks) were manually corrected using `amfbrowser`. Scale bar is given in micrometers.

**Table 1.** CNN1 performance assessed on the test set. M+: colonised root sections; M− : non-colonised root sections; Other: background and non-root sections (dusts, air bubbles, etc.).

| Network | Data augmentation | Accuracy (%) |  |  | Sensitivity (%) |  |  | Specificity (%) |  |  |
| --- | --- | --- | --- | --- | --- | --- | --- | --- | --- | --- |
|  |  | M+ | M- | Other | M+ | M- | Other | M+ | M- | Other |
| CNN1v1 | no | 98 | 96 | 98 | 94 | 99 | 95 | 100 | 95 | 100 |
| CNN1v2 | yes | 94 | 91 | 97 | 86 | 86 | 96 | 98 | 92 | 98 |

**Table 2.** CNN2 performance assessed on the test set. A: arbuscules; V: vesicles; IH: intraradical hyphae.

| Network | Data augmentation | Accuracy (%) |  |  | Sensitivity (%) |  |  | Specificity (%) |  |  |
| --- | --- | --- | --- | --- | --- | --- | --- | --- | --- | --- |
|  |  | A | V | IH | A | V | IH | A | V | IH |
| CNN2v1 | no | 94 | 95 | 93 | 94 | 97 | 94 | 94 | 89 | 88 |
| CNN2v2 | yes | 77 | 88 | 77 | 72 | 86 | 76 | 91 | 91 | 80 |

CNN2 classifiers accurately identified intraradical hyphal structures on high-magnification images **(Fig. 2b, Table 2, Fig. S4)**. The overall CNN2 performance was 94%, but dropped to 81% when data augmentation was active **(Table 2)**, suggesting this method did not improve CNN2 generalisation.

To gain more insights into AMFinder performance, we generated CNN1 confusion matrices and extracted representative sets of mispredicted tiles **(Fig. S4, S5)**. The most frequent type of CNN1 misprediction consisted of colonised root sections’ (M+) that were incorrectly identified as ‘non-colonised root sections’ (M−) **(Fig. S4a, b)**. These tiles contained low-contrasted fungal structures **(Fig. S4a, c)**. The second source of mispredictions was due to extraradical hyphae **(Fig. S4a, c)**. We then repeated the analysis with CNN2 classifiers **(Fig. S5)**. As implied by the sensitivity and specificity results **(Table 2)**, most CNN2v1 mispredictions consisted in false positives, such as vesicles or intraradical hyphae predicted in tiles not containing these structures, and confusion between arbuscules and vesicles **(Fig. S5a, c)**. CNN2 training on augmented data (CNN2v2) triggered the same mispredictions, but also increased the proportion of false negatives (non-detected structures), in particular truncated arbuscule or vesicle shapes occurring at tile edges, and low-contrasted intraradical hyphae **(Fig. S5b, c)**.

CNN1 consistently labels fungal colonisation of *N. benthamiana* roots irrespective of the image resolution and can annotate highly dissimilar image datasets, suggesting it may be compatible with a wide range of acquisition devices and staining techniques. CNN2 enables a detailed analysis of fungal hyphal structures, suggesting it may be used to monitor intraradical hyphal structure abundance within host roots. The AMFinder graphical interface allows users to inspect computer predictions and correct mispredictions to work around the limitations of tile-based image segmentation. Thus, AMFinder can robustly identify AM fungal colonisation and intraradical structures.

### AMFinder performs consistently on multiple host model species

A wide range of plants is used in endomycorrhiza research including legumes and monocot species with various root size and morphology. We assessed the suitability of AMFinder pre-trained models trained on *N. benthamiana* root images to predict AM fungal colonisation and intraradical hyphal structures on colonised root images from *Medicago truncatula* ecotype R108 **(Fig. 3a)**, *Lotus japonicus* cv. Gifu **(Fig. 3b)**, and *Oryza sativa* cv. Nipponbare **(Fig. 3c)**. The image contrast of ClearSee-treated *L. japonicus* and *M. truncatula* roots was similar to *N. benthamiana*. Conversely, large lateral roots of *O. sativa* showed higher staining background and were more challenging to destain **(Fig. 3c)**. AMFinder correctly identified roots and background in the tested images, with colonised and non-colonised root areas being accurately resolved **(Fig. 3a-c)**, including in cases where colonisation was restricted to inner cortical cell files **(Fig. 3b)**. Overall, CNN1 accuracy reached 97% **(Fig. 3, Table S5)**. Two mispredicted tiles were manually edited with `amfbrowser` **(Fig. 3b)**. We then used pre-trained CNN2 to label intraradical hyphal structures were recognised **(Fig. 3a-c)**. CNN2 correctly identified arbuscules, vesicles, and intraradical hyphae within roots of the three host plants, reaching an overall accuracy of 90% (excluding hyphopodia) **(Fig. 3, Table S6)**. Together, these findings suggest AMFinder is compatible with multiple host plants. Roots with high staining background, such as *O. sativa* large lateral roots, may require prolonged destaining for optimal analysis, or specific refinement of AMFinder pre-trained models.

**Figure 3.**
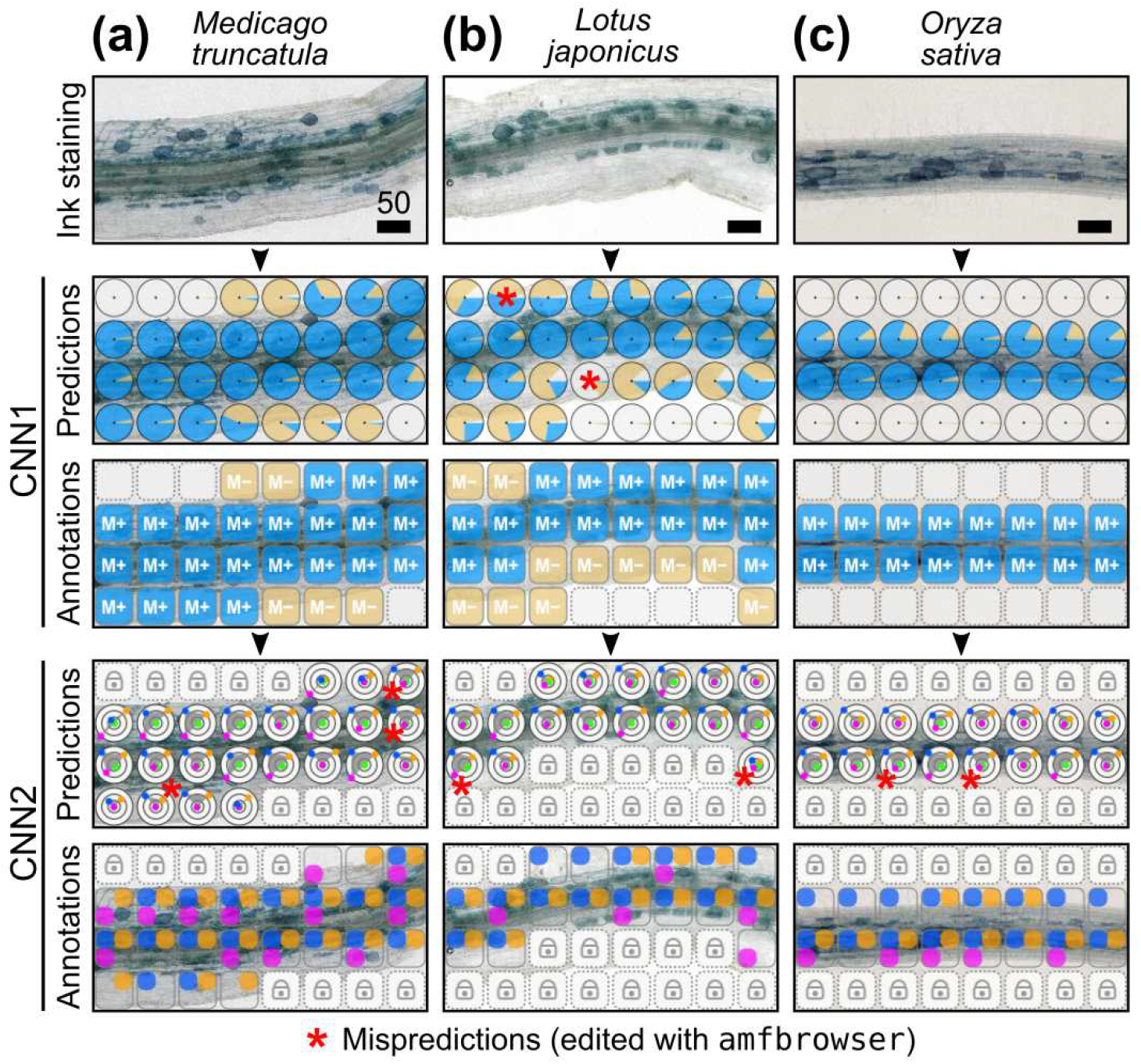
AMFinder accurately identifies AM fungal colonisation and intraradical structures in roots of various host model species. **(a-c)** Roots from *Medicago truncatula* **(a)**, *Lotus japonicus* **(b)**, and *Oryza sativa* **(c)** were stained with ink and cleared with ClearSee prior to analysis with AMFinder. `amf` mispredictions (red asterisks) were manually edited with `amfbrowser`. Scale bar is given in micrometers.

### AMFinder is compatible with multiple *Glomeraceae* species

More than a hundred AM fungal species have been reported (Chen *et al.*, 2018). To determine whether AMFinder pre-trained models are compatible with other fungal mycobionts, we analysed roots from *N. benthamiana* plants colonised with different *Glomeraceae* species **(Fig. 4)**. Colonised and non-colonised root sections were labelled by CNN1 with an overall accuracy of 97% **(Fig. 4, Table S7)**. Mispredictions (10 tiles among 160) were manually edited with `amfbrowser` **(Fig. 4)**. Then, colonised root sections were analysed with CNN2. All colonised root sections contained arbuscules and intraradical hyphae. Vesicles were abundant in sections colonised with either *Claroideoglomus claroideum* **(Fig. 4a)** or *Rhizoglomus microaggregatum* **(Fig. 4b)**, while they were scarce upon colonisation by *Funneliformis spp.* **(Fig. 4c, d)**. CNN2 correctly identified intraradical hyphal structures in all four species, reaching an overall accuracy of 87% (excluding hyphopodia) **(Fig. 4, Table S8)**. In particular, CNN2 predictions mirrored the low abundance of vesicles in *Funneliformis*-colonised roots **(Fig. 4c, d)** compared to *C. claroideum* **(Fig. 4a)** and *R. microaggregatum* **(Fig. 4b)**. Thus, AMFinder trained models are compatible with AM fungal species forming similarly-shaped intraradical hyphal structures.

**Figure 4.**
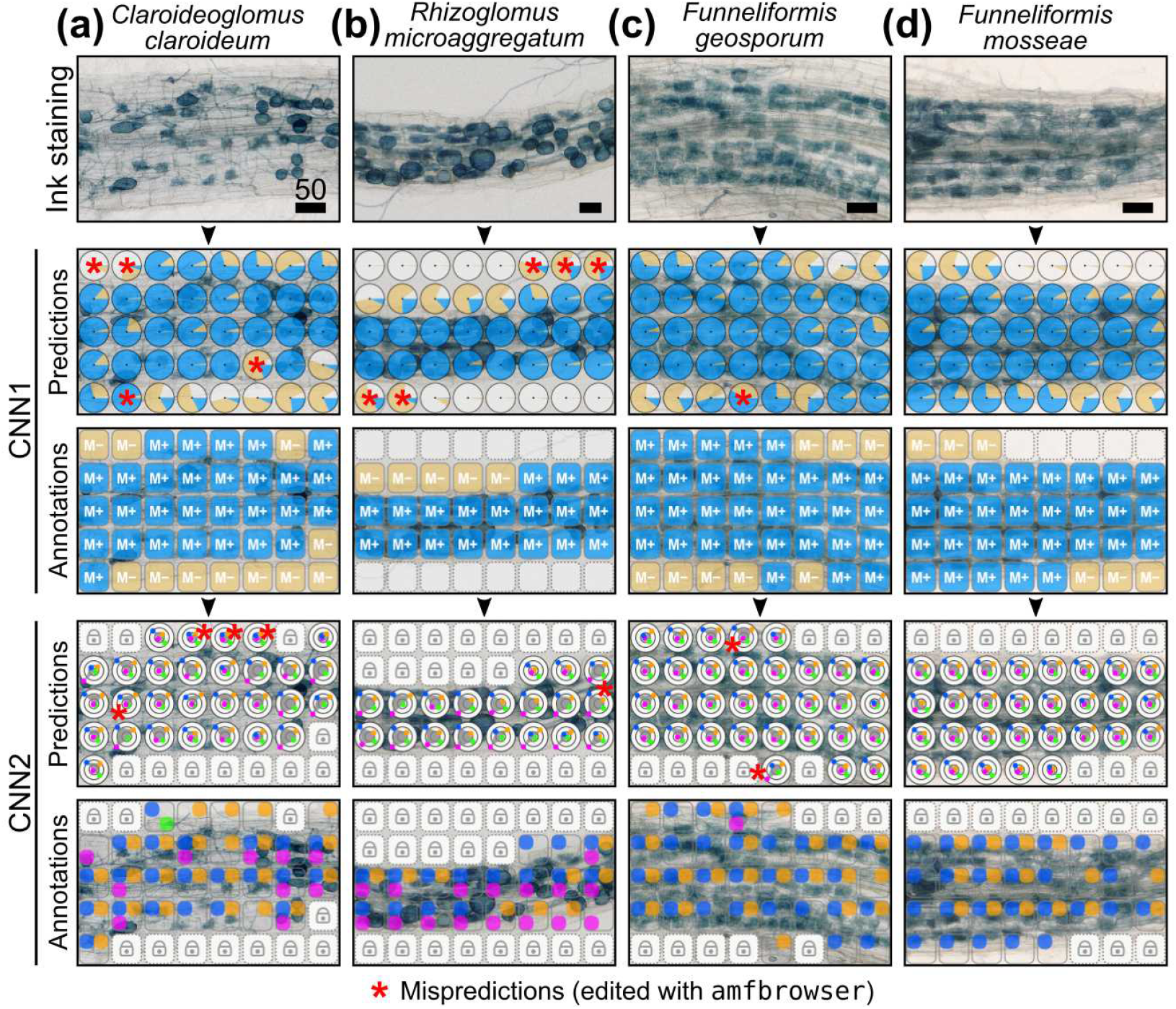
AMFinder accurately identifies colonisation and intraradical hyphal structures formed by different *Glomeraceae* species. **(a-d)** Roots from *Nicotiana benthamiana* plants growing in presence of either *Claroideoglomus claroideum* **(a)**, *Rhizoglomus microaggregatum* **(b)**, *Funneliformis geosporum* **(c)**, or *Funneliformis mosseae* **(d)**, were stained with ink and cleared with ClearSee prior to analysis with AMFinder. `amf` mispredictions (red asterisks) were manually edited with `amfbrowser`. Scale bar is given in micrometers.

### AMFinder enables *in silico* quantification of AM fungal colonisation dynamics

We next investigated whether AMFinder could be used to reliably quantify AM colonisation changes of plant roots over time. To that end, we assessed AM fungal colonisation extent on *N. benthamiana* roots harvested after a 4- or 6-week co-cultivation with *R. irregularis* **(Fig. 5)**. First, we monitored the accumulation of transcripts encoding a *N. benthamiana* ortholog of the mycorrhiza-responsive gene *MtBCP1b* (Parádi *et al.*, 2010) **(Fig. 5a)** and quantified fungal biomass by monitoring *R. irregularis EF1α* transcript levels **(Fig. 5b)**. Both methods showed a significant, two- to three-fold increase in fungal content at 6 weeks post-inoculation (wpi) compared to 4 wpi **(Fig. 5a, b)**. Then, using the grid-line intersect method (Giovannetti & Mosse, 1980) we studied the colonisation extent within randomly sampled root fragments **(Fig. 5c)**. Consistent with the molecular analysis, more colonisation was observed at 6 wpi. The analysis of the same samples with AMFinder gave similar results **(Fig. 5d)**. We then tested AMFinder’s ability to predict fungal colonisation on low-resolution flatbed scanner pictures of whole root systems **(Fig. 5e)**. Consistent with random sampling and molecular data, AM fungal colonisation levels were significantly higher at 6 wpi, although the colonisation extent values were lower than those obtained through random sampling **(Fig. 5e)**. We generated root system maps to gain insights on the distribution of colonised areas **(Fig. 5e)**. We found that while some root sections were entirely colonised, other areas were devoid of colonisation, in particular at 4 weeks post inoculation **(Fig. 5e)**, suggesting that careful mixing of root fragments is required for quantification based on a fraction of the root systems. We showcased the usefulness of CNN2 by quantifying intraradical hyphal structures on whole-root systems harvested at 2, 3 and 4 wpi **(Fig. 5f)**. Together, these results demonstrate that AMFinder allows for *in silico* quantification of AM fungal colonisation of plant roots over time including in whole root systems.

**Figure 5.**
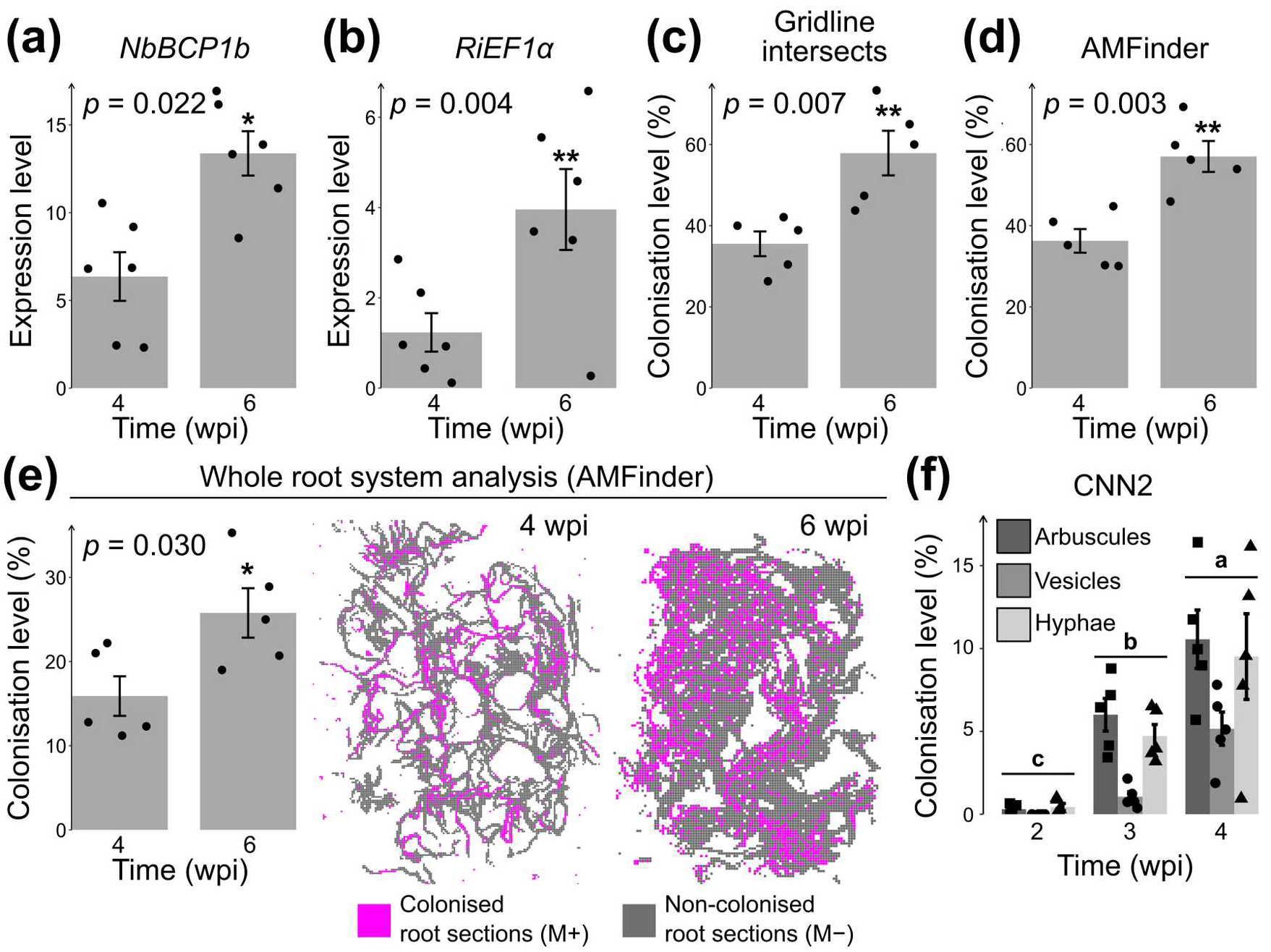
Computer vision enables quantification of AM fungal colonization on whole root systems. **(a-f)** *Nicotiana benthamiana* plants were inoculated with *Rhizophagus irregularis*. Colonisation levels were determined 4 and 6 weeks post-inoculation (wpi). **(a-b)** Quantification of transcripts corresponding to the *N. benthamiana* mycorrhizal marker gene *Blue Copper Protein 1* (*BCP1*) **(a)** and the *R. irregularis Elongation Factor 1α* (*EF1α*) **(b)**. Data are expressed relative to both *N. benthamiana L23* and *F-box* reference genes. **(c-d)** Quantification of AM fungal colonization on ink-stained root pictures using the gridline intersect method **(c)**, or AMFinder **(d)**. **(e)** Quantification of AM fungal colonisation on whole root systems using AMFinder, and representative images of computer-generated maps featuring colonized (M+, magenta) and non-colonized (M−, grey) root areas at 4 and 6 wpi. **(f)** Intraradical hyphal structures were quantified on whole-root systems at 2, 3, and 4 wpi using CNN2. Dots correspond to biological replicates. Bars represent standard error. Statistical significance was assessed using Student’s *t*-test (*: *p* < 0.05; **: *p* < 0.01).

### AMFinder accurately quantifies colonisation levels of *ram1*, *str*, and *smax1* mutants

To determine whether AMFinder can identify altered colonisation levels in plant mutants, we quantified colonisation levels of *M. truncatula* roots, either wild-type or carrying the mutations *ram1-1* (Gobbato *et al.*, 2012) and *str* (Zhang *et al.*, 2010) **(Fig. 6a-c)**. Consistent with the phenotypes reported for these mutants, we found that the extent of root colonisation by *irregularis* was reduced in *ram-1-1* and *str* backgrounds compared to wild-type **(Fig. 6a)**. Similarly, both mutants showed reduced amounts of arbuscules, vesicles, and intraradical hyphae compared to wild-type, and arbuscules and vesicles were rare in the str mutant **(Fig. 6b, c)**. Microscopic inspection of the colonised roots showed stunted arbuscules in the *ram1-1* and *str* mutant backgrounds compared to wild-type, with a more severe reduction in arbuscule branching in *str* **(Fig. 6c)**. Next, we investigated whether AMFinder can detect an increased colonisation. We assessed the extent of *R. irregularis* colonisation of *O. sativa* roots either wild-type or carrying a mutation that abolishes expression of the suppressor of AM symbiosis SMAX1 (Choi *et al.*, 2020). We found that colonisation level was significantly increased in smax1 mutant compared to wild-type **(Fig. 6d, f)**. In addition, CNN2 analysis showed that all types of intraradical hyphal structures were more abundant in smax1 roots **(Fig. 6e)**. Thus, AMfinder accurately detects the AM fungal colonisation phenotypes of well-established plant mutants, suggesting it can adapt to multiple host genetic backgrounds.

**Figure 6.**
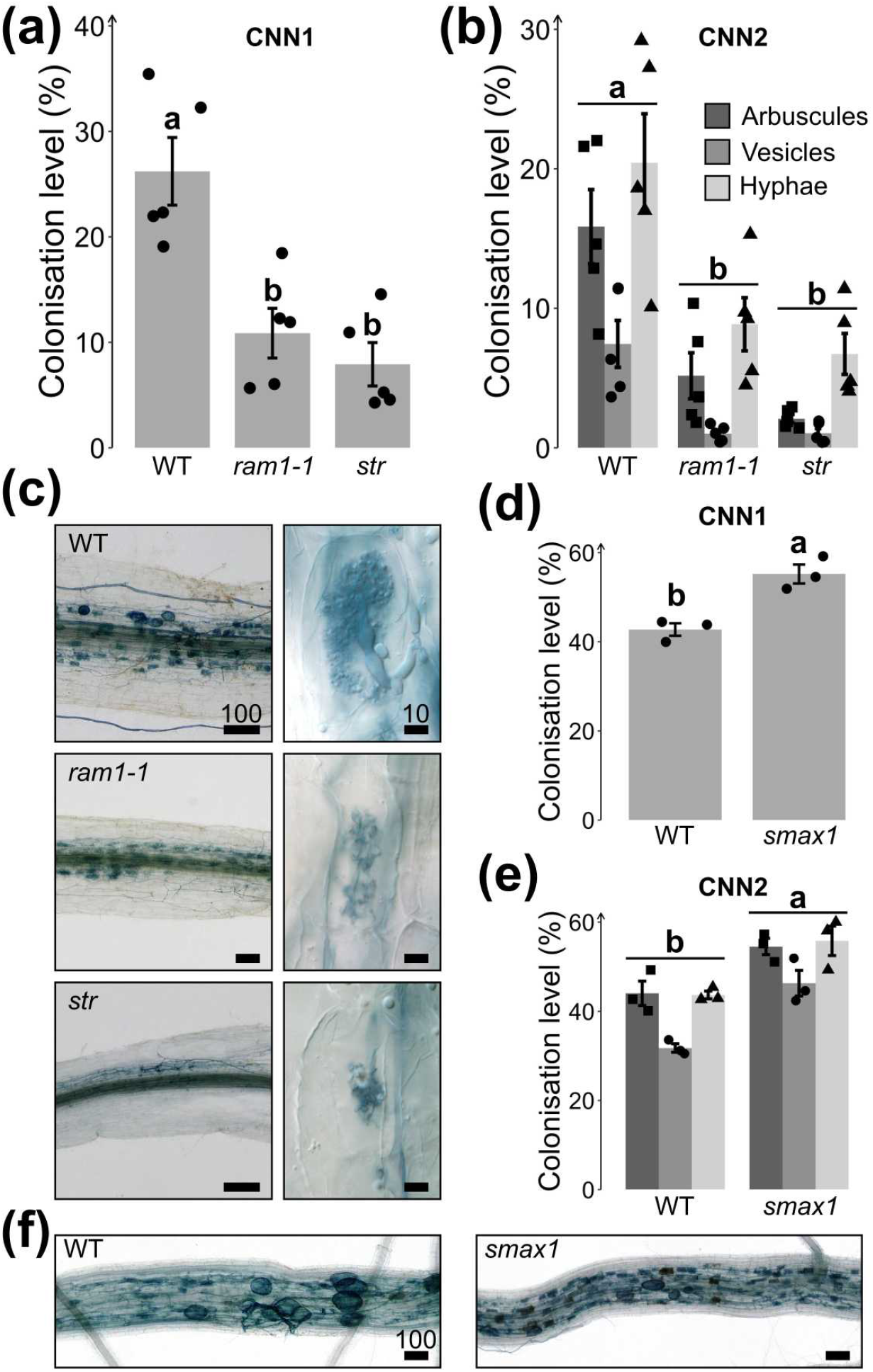
AMFinder accurately quantifies altered colonisation levels of *ram1*, *str*, and *smax1* mutants. **(a-c)** *Medicago truncatula* A17 (WT), *ram1-1*, and *str* were grown on sand for 6 weeks in presence of *Rhizophagus irregularis*. Barplots show overall colonisation level obtained using CNN1 and a detailed analysis of root content in arbuscules, vesicles, and intraradical hyphae obtained by CNN2 analysis **(b)**. Representative images of colonised roots and arbuscule shape **(c)**. **(d-f)***Oryza sativa* wild-type and *smax1* were grown on sand for 6 weeks in presence of *R. irregularis*. Barplots show overall colonisation levels **(d)** and quantification of intraradical hyphal structures **(e)**. **(f)** Representative images of colonised large lateral roots. Statistical significance was assessed using ANOVA and Tukey’s HSD (P < 0.05). Letters indicate statistical groupings. Bars represent the standard error. Scale bars are given in micrometers.

## Discussion

We developed the software AMFinder, which uses two convolutional neural networks to annotate and quantify AM fungi in plant roots. AMFinder performs consistently well on root images of several model plant and fungal species used in endosymbiosis research. AMFinder-mediated quantification of AM fungal colonisation gives similar results to those obtained using current standard counting methods. We further show that AMFinder can process whole root systems using low-resolution flatbed scans obtained from an optimised ink-staining protocol which relies on ClearSee as a contrast enhancer. We illustrate the usefulness of this approach to study fungal colonisation dynamics over time in wild-type and mutant plants.

AMFinder enables computer-assisted quantification of AM fungal colonisation. A pioneering attempt to automate the quantification of AM fungal colonisation relied on pixel-based image segmentation to determine projected root and fungal surface areas using the proprietary software WinRHIZO developed by Regent Instruments (Kokkoris *et al.*, 2019). Similar to AMFinder, this method enabled whole-root-system analyses and showed that quantification based on gridline intersects was generally overestimated (Kokkoris *et al.*, 2019). By contrast with pixel-based methods, CNNs are trainable and can thus adapt to a wide range of images, including environmental samples colonised with both AM and non-AM fungal species. Furthermore, CNNs further enable the quantification of different types of intraradical hyphal structures, which are important parameters of plant mutant phenotypes. Thus, deep learning ensures AMFinder versatility and enables a more detailed analysis of mycorrhizal phenotypes.

AMFinder’s design adequately addresses limitations arising from its computer vision approach while still enabling a low- to medium-throughput workflow. Specifically, we implemented a semi-automatic pipeline that requires user supervision of computer predictions. High-throughput AMFinder analyses, such as large-scale field experiments, would require the entire prediction pipeline to be fully automatic, including the conversion to annotations. However, input image parameters can influence pre-trained model accuracy and may require user adjustments. Automatic analyses assuming image suitability without quality control may overestimate CNN model accuracy. Besides, CNN2 predictions on mispredicted M+ tiles without intraradical hyphal structures have not been investigated in this study. Besides, this AMFinder implementation does not discriminate between multiple root types. As a result, image data from experiments relying on crude inoculum (Habte & Byappanhalli, 1998) or nurse root systems such as chives (Demchenko *et al.*, 2004) as an inoculation method may pose problems when contaminating root fragments remain in root images. To that end, AMFinder offers manual curation of predictions to avoid systematic introduction of errors.

We used an uncertainty sampling-like bootstrap method, wherein CNNs are trained on a small amount of manually annotated images, and then the user is prompted to label further training images which the model is unsure about, for rapid creation of a large training dataset. Despite misannotations in the resulting training dataset, AMFinder trained networks showed high accuracy on a large set of images featuring different plant and fungal species, confirming the suitability of this strategy. Further improvements could possibly be achieved using an active learning (Shen *et al.*, 2018).

We trained and tested AMFinder on ink-vinegar stained *N. benthamiana* roots. Ink-vinegar is an inexpensive, non-toxic fungal staining method compatible with various plant and mycobiont species (Vierheilig *et al.*, 1998). Thus, pre-trained CNNs generated from ink-stained roots ensure immediate workability of AMFinder for most endosymbiosis host systems without the need to generate manually-annotated training datasets. Besides, we showed that data augmentation allows for accurate labelling of images with altered hue, intensity and background colours. If needed, AMFinder can be trained with datasets obtained using other dyes and fluorophores for fungal staining (Vierheilig *et al.*, 2005) or for the annotation of other tissues colonised by fungi such as liverwort thalli (Ligrone *et al.*, 2007; Carella & Schornack, 2018; Kobae *et al.*, 2019). Computer-intensive computations required for *ab initio* training can be avoided by refining the existing pre-trained networks. Thus, AMFinder is highly versatile and can be adapted for studying many aspects of fungal colonisation; it may also be of interest to researchers of pathogenic fungi.

AMFinder sensitivity is best on vesicles, likely because AM fungal vesicles are fairly invariant, globular-shaped structures surrounded by a thick, multilayered wall (Jabaji-Hare *et al.*, 1990) that result in high contrast signals within the surrounding plant tissues. By contrast, the arbuscular shape is more diverse, with branching extent and cell volume occupancy increasing during the initial development stages (Toth & Miller, 1984) and a size that is ultimately defined by host cell size. Intraradical hyphae show different diameters, orientations, and staining intensities, and occasionally overlay other intraradical structures. Besides, the limited pixel information of a single tile may not always discriminate between intraradical and extraradical hyphae. An approach using information from a wider area of the original image, rather than treating each tile in isolation, may help to address this issue. In particular, it would be interesting to apply deep learning image segmentation techniques (Ghosh *et al.*, 2019) to this problem, as researchers have often found success with this approach in other types of biological imaging. Another possible issue is that convolutional neural networks do not retain relative spatial information (Patrick *et al.*, 2019). Solutions to overcome this limitation include the combination of convolutional neural networks and multi-layer perceptrons (Haldekar *et al.*, 2017), and capsule networks (CapsNets) (Sabour *et al.*, 2017; Patrick *et al.*, 2019). Future work will explore the usefulness of such approaches to achieve even higher prediction accuracy.

Obtaining contrasted fungal structures within root tissues is pivotal for accurate AMFinder predictions. The first report of ink-vinegar staining of AM fungi suggests that black and blue inks allow for high-contrast images in at least four plant species (Vierheilig *et al.*, 1998). Background destaining in tap water with few vinegar droplets requires at least 20 min incubation and is only effective against excess ink (Vierheilig *et al.*, 1998). By contrast, ClearSee treatment works in seconds and allows for both destaining and clearing (Kurihara *et al.*, 2015). Such a feature is of particular interest for thick or pigmented roots, and soil samples. Also, ClearSee preserves fluorescence (Kurihara *et al.*, 2015) and is thus compatible with immunohistochemical fungal labelling techniques such as wheat germ agglutinin-fluorophore conjugates (Bonfante-Fasolo *et al.*, 1990).

AMFinder can improve the robustness and reproducibility of AM fungal quantification. In the gridline-intersect method, gridlines have been primarily used as guides for the systematic selection of observation points (Giovannetti & Mosse, 1980), and the distance between adjacent lines has been studied to estimate the total root length, but not to improve quantification accuracy (Newman, 1966; Marsh, 1971; Giovannetti & Mosse, 1980). As a result, a low number of root fragments is considered prejudicial to quantification accuracy (Giovannetti & Mosse, 1980). Also, the shape of the area surrounding the grid-root intersection used for visual scoring has not been formally described and may account for variations between experimenters. By contrast, AMFinder analyses well-defined tiles, and tile size can adjust to image resolution without impairing prediction accuracy. Intraradical hyphal structures cannot be identified from flatbed scans due to limited resolution. However, machine learning-based algorithms have been recently developed to achieve data-driven image super-resolution (Park *et al.*, 2003; Wang *et al.*, 2019). By contrast with standard image interpolation techniques, super-resolution algorithms predict missing details by learning common patterns from training datasets. Whether such algorithms can enable a detailed analysis of AM fungal hyphal structures from flatbed scans could be explored in future AMFinder developments.

## Conclusions

We have demonstrated that AMFinder adapts to different plant and fungal species, fungal staining methods and acquisition devices. Its design ensures user control over the annotation process and facilitates data visualisation in the context of the root images. As such, it improves documentation and reproducibility of AM fungal colonisation analyses.

## Acknowledgements

We thank Raymond Wightman (SLCU, Cambridge) for assistance with digital microscopy. We are grateful to Johannes Stuttmann (Martin-Luther-Universität Halle-Wittenberg, Germany) for critical comments on the manuscript. We thank Colleen Drapek (SLCU, Cambridge) for providing us with *Funneliformis mosseae* inoculum. This work was funded by the Gatsby Charitable Foundation (GAT3395/GLD), by the European Research Council (ERC-2014-STG, H2020, and 637537), and by the Royal Society (UF110073 and UF160413). C.T. is supported by a Junior Research Fellowship at Gonville & Caius College, Cambridge, and also acknowledges support by the STFC consolidated grant ST/P000681/1.

## Author Contribution

E.E. developed the software, conceived the experimental strategy, conducted experiments, acquired data, analysed data and wrote the manuscript. C.T. developed the software and analysed data. A. M. conceived the original idea, conducted experiments and acquired data. L.S., T.Y. and A.G. conducted experiments and acquired data. E.S., C.Q. and J.S. contributed material. S.S. acquired funding, conceived the experimental strategy, analysed data and wrote the manuscript. All authors have read and approved the manuscript.

## Data Availability

AMFinder is released under the terms of the open-source MIT license (https://opensource.org/licenses/MIT) allowing unrestricted usage. Source code, pre-trained models and detailed installation instructions are available on AMFinder GitHub webpage (https://github.com/SchornacklabSLCU/amfinder.git). Training datasets are available upon request.

## Supporting information

Figure S1. ClearSee enhances the contrast of ink-stained roots.

Figure S2. Optimisation of flatbed scanner resolution for ink-stained root imaging.

Figure S3. Schematic representation of AMFinder ConvNet architecture.

Figure S4. Analysis of CNN1 mispredictions.

Figure S5. Analysis of CNN2 mispredictions.

Table S1. Primers used in this study.

Table S2. CNN1 training dataset.

Table S3. CNN2 training dataset.

Table S4. Test dataset.

Table S5. CNN1 performance on various host plants.

Table S6. CNN2 performance on various host plants.

Table S7. CNN1 performance on various AM fungi.

Table S8. CNN2 performance on various AM fungi.

