## Supplementary Information for "Deep learning-based quantification of arbuscular mycorrhizal fungi in plant roots"

##### Supporting figures

**Figure S1.** ClearSee enhances the contrast of ink-stained roots. 2

**Figure S2.** Optimisation of flatbed scanner resolution for ink-stained root imaging. 2

**Figure S3.** Schematic representation of AMFinder ConvNet architecture. 3

**Figure S4.** Analysis of CNN1 mispredictions. 4

**Figure S5.** Analysis of CNN2 mispredictions. 5

##### Supporting tables

**Table S1.** Primers used in this study. 6

**Table S2.** CNN1 training dataset. 6

**Table S3.** CNN2 training dataset. 7

**Table S4.** Test dataset. 8

**Table S5.** CNN1 performance on various host plants. 8

**Table S6.** CNN2 performance on various host plants. 8

**Table S7.** CNN1 performance on various AM fungi. 9

**Table S8.** CNN2 performance on various AM fungi. 9

### Supporting figures

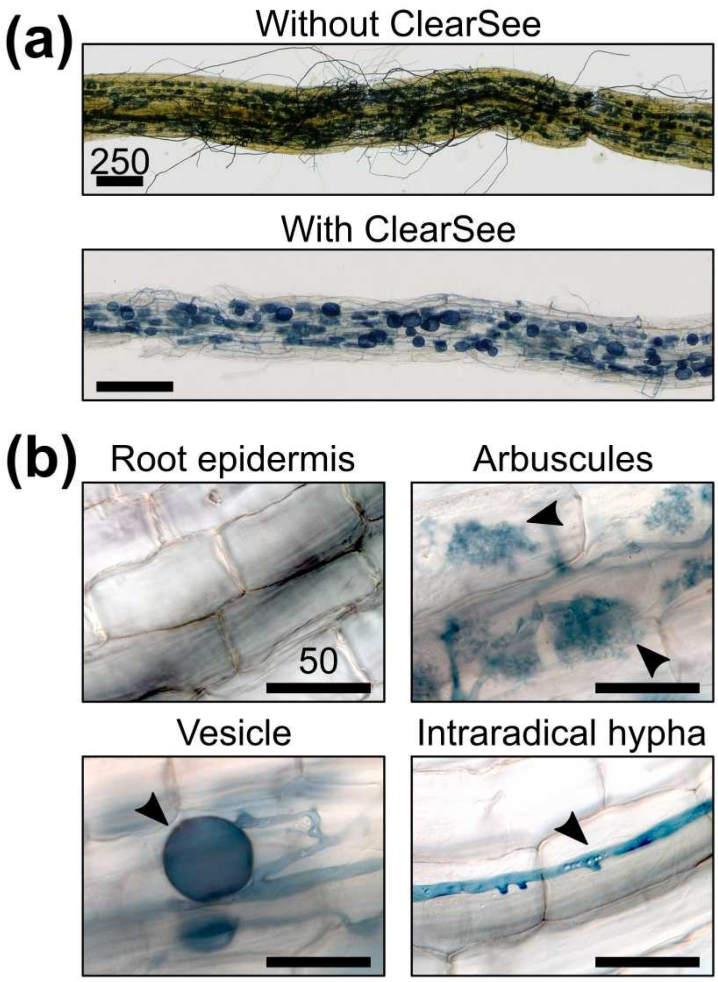


**Figure S1. ClearSee enhances the contrast of ink-stained roots. (a)** Representative pictures of *Nicotiana benthamiana* roots without (upper pane) or with (lower pane) clearing with ClearSee. **(b)** Magnified views of *N. benthamiana* root epidermis, and *Rhizophagus irregularis* intraradical structures (arbuscules, vesicles and intraradical hyphae) from ClearSee-treated ink-stained samples. Scale bar is given in micrometers.

**Figure S2. Optimisation of flatbed scanner resolution for ink-stained root imaging.** Representative pictures of an ink-stained *Nicotiana benthamiana* root acquired at scanner resolutions ranging from 800 to 12800 dots per inch (dpi). The bottom right pane shows the same area acquired using a microscope. The resolution of the red-framed pane (3200 dpi) was used to generate the pictures analysed in this study. Scale bar is 100 µm.

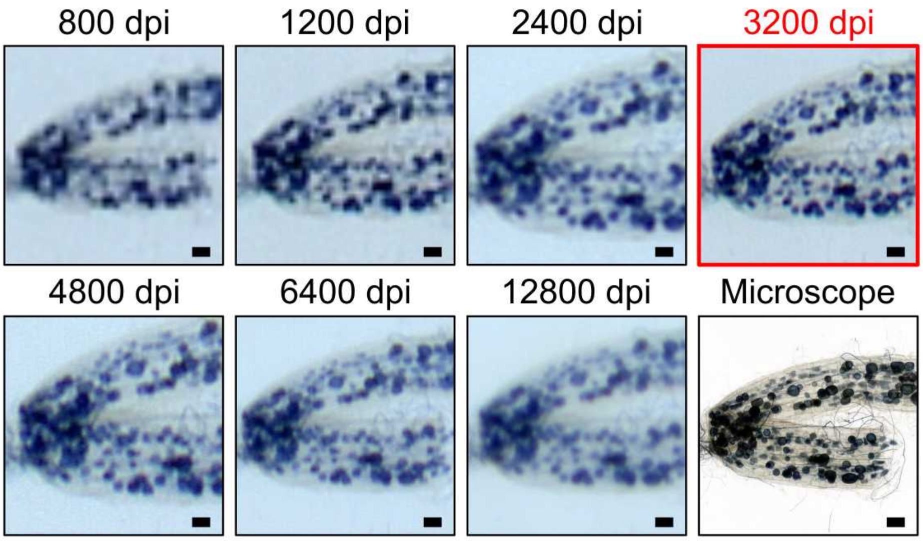

**Figure S3. Schematic representation of AMFinder ConvNet architecture. (a)** Input images (tiles) are resized to 126×126 pixels and processed by four blocks of 3×3 convolutions (conv) followed by maximum pooling (maxpool), resulting in size reduction from 126×126 pixels to 5×5 pixels. **(b)** ConvNet 1 (CNN1) uses three fully connected layers (fc). The last layer predicts one out of three classes (colonized, non-colonized, background) using a normalized exponential (softmax) as the activation function. There are approximately 1.5 million trainable parameters. **(c)** ConvNet 2 (CNN2) relies on four independent sets of fully connected layers. Each set is organized as in **b**, except that the last layer predicts a single AM fungal structure (either arbuscules, vesicles, or intraradical hyphae) using a sigmoid activation function. There are approximately 4 million trainable parameters.

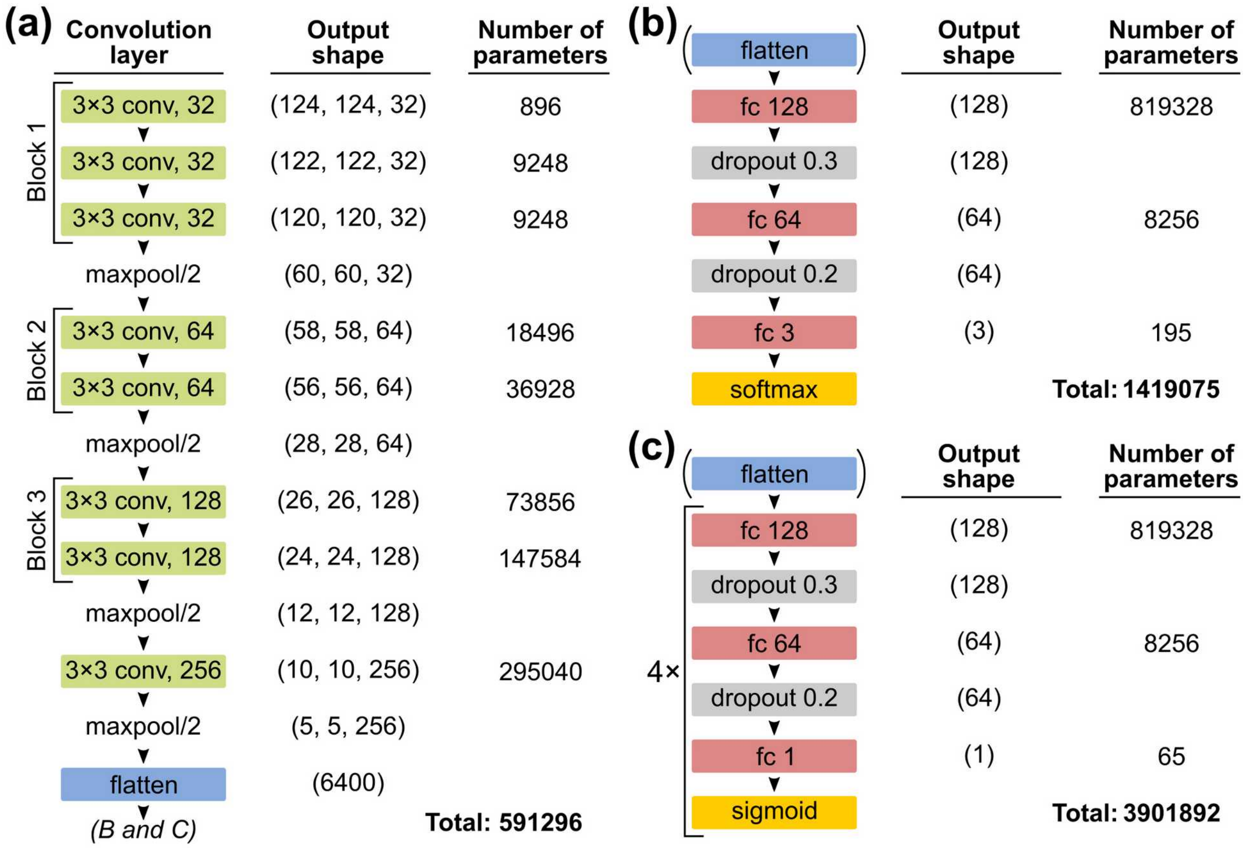

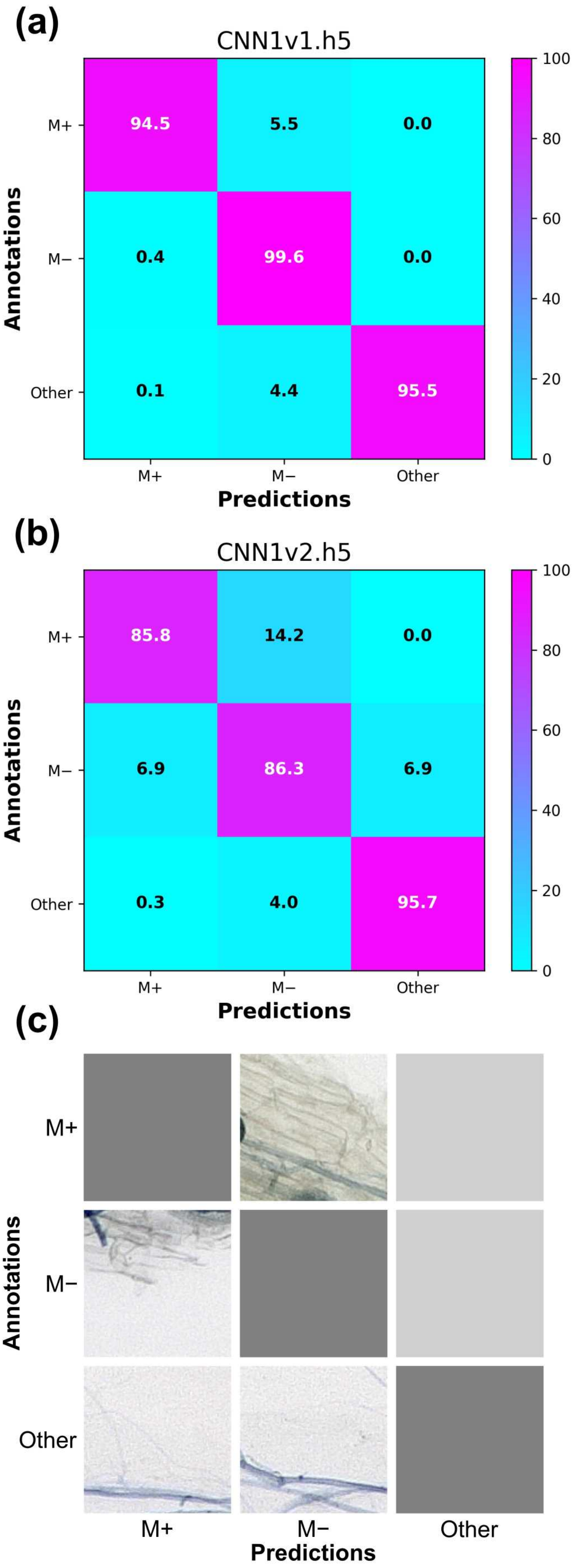


**Figure S4. Analysis of CNN1 mispredictions.** CNN1 was used to identify colonised root sections on training images. Computer probabilities were then compared with human annotations to compute row-normalized confusion matrices and extract subsets of mispredicted tiles. **(a)** Confusion matrix of CNN1v1. **(b)** Confusion matrix of CNN1v2. **(c)** Representative examples of CNN1v1 mispredicted tiles, displayed in a similar way to confusion matrices. Light grey squares reflect that no more tiles are available in a given misprediction group.

**Figure S5. Analysis of CNN2 mispredictions.** CNN2 was used to identify colonised root sections on training images. Computer probabilities were then compared with human annotations to compute row-normalized confusion matrices and extract subsets of mispredicted tiles. **(a)** Confusion matrix of CNN2v1. **(b)** Confusion matrix of CNN2v2. **(c)** Representative examples of CNN2v1 mispredicted tiles, displayed in a similar way toconfusion matrices. A: arbuscules; V: vesicles: I: intraradical hyphae.

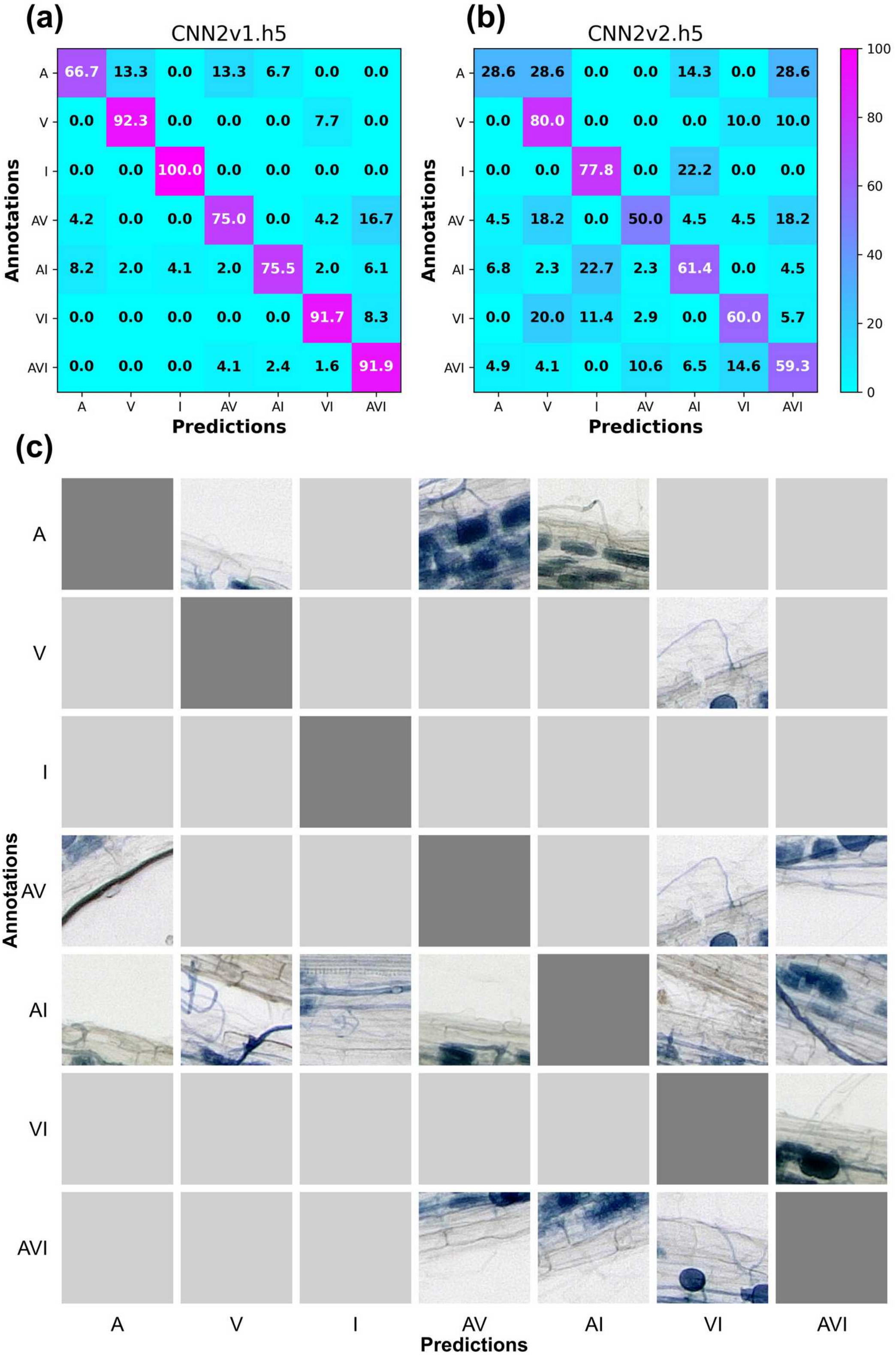

### Supporting tables

**Table S1. Primers used in this study.**

| Primer name | Sequence (5′ → 3′) |
|---|---|
| RiEF1a_F | TGTTGCTTTCGTCCCAATATC |
| RiEF1a_R | GGTTTATCGGTAGGTCGAG |
| NbBCP1b_F | TCGAAGATGGAGTTCCGGCA |
| NbBCP1b_R | AATCAACATCAATGGTCCAGCC |

**Table S2. CNN1 training dataset.** M+: tiles labelled as colonised root sections; M−: tiles labelled as non-colonised root sections; Other: any other tile (background, debris, air bubbles, etc.); Discarded: number of background tiles randomly removed during CNN1 training. px: pixels.

| # | Image name | Acquisition device | Image size (px) | Tile size (px) | M+ | M− | Other | Discarded |
|---|---|---|---|---|---|---|---|---|
| 1 | Background_1.jpg | Flatbed scanner | 3608 x 860 | 40 | 0 | 0 | 1890 | 1673 |
| 2 | Background_2.jpg | Flatbed scanner | 3496 x 964 | 40 | 0 | 0 | 2088 | 1848 |
| 3 | Flatbed_scan_Nbenth_1.jpg | Flatbed scanner | 6272 x 3168 | 40 | 44 | 2335 | 9945 | 8764 |
| 4 | Flatbed_scan_Nbenth_2.jpg | Flatbed scanner | 6272 x 3168 | 40 | 38 | 2240 | 10046 | 8846 |
| 5 | Flatbed_scan_Nbenth_3.jpg | Flatbed scanner | 10454 x 8139 | 40 | 1059 | 8934 | 42986 | 37788 |
| 6 | Keyence_Nbenth_myc_1.jpg | Digital microscope | 20004 x 18208 | 256 | 1014 | 219 | 4305 | 3792 |
| 7 | Keyence_Nbenth_myc_2.jpg | Digital microscope | 18487 x 15351 | 256 | 868 | 302 | 3078 | 2714 |
| 8 | Keyence_Nbenth_myc_3.jpg | Digital microscope | 14908 x 16371 | 256 | 805 | 203 | 2646 | 2325 |
| 9 | Keyence_Nbenth_myc_4.jpg | Digital microscope | 15396 x 13385 | 256 | 846 | 236 | 2038 | 1811 |
| 10 | Keyence_Nbenth_myc_5.jpg | Digital microscope | 18020 x 14680 | 256 | 934 | 292 | 2764 | 2437 |
| 11 | Keyence_Nbenth_myc_6.jpg | Digital microscope | 15818 x 17569 | 256 | 896 | 388 | 2864 | 2540 |
| 12 | Keyence_Nbenth_myc_7.jpg | Digital microscope | 16632 x 17376 | 256 | 1096 | 230 | 2962 | 2602 |
| 13 | Keyence_Nbenth_myc_8.jpg | Digital microscope | 9560 x 3204 | 126 | 325 | 152 | 1398 | 1237 |
| 14 | Keyence_Nbenth_myc_9.jpg | Digital microscope | 9560 x 3205 | 126 | 321 | 127 | 1427 | 1249 |
| 15 | Keyence_Nbenth_myc_10.jpg | Digital microscope | 9560 x 3204 | 126 | 384 | 183 | 1308 | 1172 |
| 16 | Keyence_Nbenth_myc_11.jpg | Digital microscope | 12541 x 4044 | 126 | 405 | 140 | 2623 | 2287 |
| 17 | Keyence_Nbenth_myc_12.jpg | Digital microscope | 12541 x 4043 | 126 | 442 | 167 | 2559 | 2230 |
| 18 | Keyence_Nbenth_myc_13.jpg | Digital microscope | 12541 x 4044 | 126 | 240 | 107 | 2821 | 2471 |
| 19 | Keyence_Nbenth_myc_14.jpg | Digital microscope | 10942 x 3404 | 126 | 237 | 163 | 1922 | 1687 |
| 20 | Keyence_Nbenth_myc_15.jpg | Digital microscope | 10942 x 3403 | 126 | 425 | 217 | 1680 | 1479 |
| 21 | Keyence_Nbenth_myc_16.jpg | Digital microscope | 10942 x 3404 | 126 | 333 | 174 | 1815 | 1564 |
| 22 | Keyence_Nbenth_myc_17.jpg | Digital microscope | 14639 x 4013 | 126 | 328 | 288 | 2979 | 2629 |
| 23 | Keyence_Nbenth_myc_18.jpg | Digital microscope | 14639 x 4012 | 126 | 372 | 205 | 3019 | 2673 |
| 24 | Keyence_Nbenth_myc_19.jpg | Digital microscope | 14639 x 4013 | 126 | 431 | 296 | 2869 | 2532 |
| 25 | Keyence_Nbenth_myc_20.jpg | Digital microscope | 11049 x 5630 | 126 | 336 | 193 | 3299 | 2951 |
| 26 | Keyence_Nbenth_myc_21.jpg | Digital microscope | 11049 x 5630 | 126 | 477 | 275 | 3076 | 2706 |
| 27 | Keyence_Nbenth_myc_22.jpg | Digital microscope | 11399 x 5239 | 126 | 336 | 159 | 3195 | 2826 |
| 28 | Keyence_Nbenth_myc_23.jpg | Digital microscope | 11399 x 5238 | 126 | 582 | 204 | 2904 | 2555 |
| 29 | Keyence_Nbenth_myc_24.jpg | Digital microscope | 12152 x 4747 | 126 | 399 | 150 | 3003 | 2645 |
| 30 | Keyence_Nbenth_myc_25.jpg | Digital microscope | 12152 x 4746 | 126 | 399 | 223 | 2930 | 2580 |
| 31 | Keyence_Nbenth_myc_26.jpg | Digital microscope | 12152 x 4747 | 126 | 442 | 134 | 2976 | 2616 |
| 32 | Keyence_Nbenth_myc_27.jpg | Digital microscope | 16632 x 5792 | 126 | 550 | 519 | 4871 | 4282 |
| | | | | **Count** | **15364** | **19455** | **140286** | **123511** |

**Table S3. CNN2 training dataset.** A: tiles containing arbuscules; V: tiles containing vesicles; H: tiles containing hyphopodia; IH: tiles containing intraradical hyphae; px: pixels.

| # | Image name | Image size (px) | Tile size (px) | A | V | H | IH |
|---|---|---|---|---|---|---|---|
| 1 | Keyence_Nbenth_myc_10.jpg | 9560 x 3204 | 126 | 315 | 284 | 1 | 177 |
| 2 | Keyence_Nbenth_myc_11.jpg | 12541 x 4044 | 126 | 351 | 322 | 20 | 171 |
| 3 | Keyence_Nbenth_myc_12.jpg | 12541 x 4043 | 126 | 336 | 293 | 17 | 240 |
| 4 | Keyence_Nbenth_myc_13.jpg | 12541 x 4044 | 126 | 204 | 171 | 9 | 154 |
| 5 | Keyence_Nbenth_myc_14.jpg | 10942 x 3404 | 126 | 183 | 185 | 9 | 153 |
| 6 | Keyence_Nbenth_myc_15.jpg | 10942 x 3403 | 126 | 338 | 270 | 8 | 269 |
| 7 | Keyence_Nbenth_myc_16.jpg | 10942 x 3404 | 126 | 255 | 179 | 4 | 254 |
| 8 | Keyence_Nbenth_myc_17.jpg | 14639 x 4013 | 126 | 249 | 207 | 6 | 191 |
| 9 | Keyence_Nbenth_myc_18.jpg | 14639 x 4012 | 126 | 306 | 271 | 10 | 284 |
| 10 | Keyence_Nbenth_myc_19.jpg | 14639 x 4013 | 126 | 322 | 249 | 9 | 341 |
| 11 | Keyence_Nbenth_myc_20.jpg | 11049 x 5630 | 126 | 269 | 224 | 2 | 245 |
| 12 | Keyence_Nbenth_myc_21.jpg | 11049 x 5630 | 126 | 388 | 257 | 11 | 371 |
| 13 | Keyence_Nbenth_myc_22.jpg | 11399 x 5239 | 126 | 250 | 188 | 7 | 275 |
| 14 | Keyence_Nbenth_myc_23.jpg | 11399 x 5238 | 126 | 496 | 357 | 13 | 432 |
| 15 | Keyence_Nbenth_myc_27.jpg | 16632 x 5792 | 126 | 222 | 224 | 9 | 281 |
| 16 | Keyence_Nbenth_myc_28.jpg | 4832 x 2220 | 126 | 94 | 89 | 3 | 197 |
| 17 | Keyence_Nbenth_myc_29.jpg | 11810 x 4062 | 126 | 141 | 144 | 1 | 145 |
| 18 | Keyence_Nbenth_myc_30.jpg | 11810 x 4062 | 126 | 268 | 190 | 11 | 267 |
| 19 | Keyence_Nbenth_myc_31.jpg | 11810 x 4062 | 126 | 418 | 374 | 10 | 419 |
| 20 | Keyence_Nbenth_myc_32.jpg | 11049 x 5630 | 126 | 325 | 250 | 6 | 235 |
| 21 | Keyence_Nbenth_myc_33.jpg | 11399 x 5239 | 126 | 246 | 167 | 4 | 307 |
| 22 | Keyence_Nbenth_myc_34.jpg | 10389 x 5108 | 126 | 368 | 298 | 10 | 207 |
| 23 | Keyence_Nbenth_myc_35.jpg | 10389 x 5108 | 126 | 422 | 408 | 14 | 374 |
| 24 | Keyence_Nbenth_myc_36.jpg | 10389 x 5108 | 126 | 202 | 184 | 5 | 202 |
| 25 | Keyence_Nbenth_myc_37.jpg | 11593 x 5269 | 126 | 348 | 305 | 7 | 337 |
| 26 | Keyence_Nbenth_myc_38.jpg | 11593 x 5268 | 126 | 356 | 299 | 4 | 373 |
| 27 | Keyence_Nbenth_myc_39.jpg | 11593 x 5269 | 126 | 367 | 321 | 7 | 395 |
| 28 | Keyence_Nbenth_myc_40.jpg | 14301 x 5439 | 126 | 672 | 616 | 18 | 691 |
| 29 | Keyence_Nbenth_myc_41.jpg | 14301 x 5440 | 126 | 681 | 492 | 10 | 733 |
| 30 | Keyence_Nbenth_myc_42.jpg | 14301 x 5439 | 126 | 461 | 380 | 5 | 530 |
| 31 | Keyence_Nbenth_myc_43.jpg | 11488 x 4755 | 126 | 268 | 193 | 7 | 282 |
| 32 | Keyence_Nbenth_myc_44.jpg | 11488 x 4754 | 126 | 333 | 262 | 11 | 347 |
| 33 | Keyence_Nbenth_myc_45.jpg | 11488 x 4755 | 126 | 303 | 204 | 11 | 327 |
| 34 | Keyence_Nbenth_myc_46.jpg | 20004 x 6069 | 126 | 393 | 249 | 7 | 491 |
| 35 | Keyence_Nbenth_myc_47.jpg | 20004 x 6070 | 126 | 633 | 462 | 30 | 867 |
| 36 | Keyence_Nbenth_myc_48.jpg | 20004 x 6069 | 126 | 442 | 271 | 14 | 609 |
| 37 | Keyence_Nbenth_myc_49.jpg | 18487 x 5117 | 126 | 353 | 292 | 30 | 683 |
| 38 | Keyence_Nbenth_myc_50.jpg | 18487 x 5117 | 126 | 427 | 428 | 3 | 745 |
| 39 | Keyence_Nbenth_myc_51.jpg | 18487 x 5117 | 126 | 402 | 331 | 15 | 781 |
| 40 | Keyence_Nbenth_myc_52.jpg | 14908 x 5457 | 126 | 425 | 326 | 12 | 720 |
| 41 | Keyence_Nbenth_myc_53.jpg | 14908 x 5457 | 126 | 516 | 452 | 12 | 688 |
| 42 | Keyence_Nbenth_myc_54.jpg | 14908 x 5457 | 126 | 322 | 261 | 5 | 521 |
| 43 | Keyence_Nbenth_myc_55.jpg | 15396 x 4462 | 126 | 275 | 244 | 0 | 542 |
| 44 | Keyence_Nbenth_myc_56.jpg | 15396 x 4461 | 126 | 418 | 394 | 0 | 519 |
| 45 | Keyence_Nbenth_myc_57.jpg | 15396 x 4462 | 126 | 570 | 418 | 1 | 827 |
| 46 | Keyence_Nbenth_myc_58.jpg | 18020 x 4893 | 126 | 368 | 325 | 11 | 613 |
| 47 | Keyence_Nbenth_myc_59.jpg | 18020 x 4894 | 126 | 511 | 343 | 6 | 879 |
| 48 | Keyence_Nbenth_myc_60.jpg | 18020 x 4893 | 126 | 398 | 274 | 11 | 768 |
| 49 | Keyence_Nbenth_myc_61.jpg | 15818 x 5856 | 126 | 390 | 270 | 0 | 531 |
| 50 | Keyence_Nbenth_myc_62.jpg | 15818 x 5857 | 126 | 626 | 540 | 28 | 1072 |
| 51 | Keyence_Nbenth_myc_63.jpg | 15818 x 5856 | 126 | 312 | 249 | 4 | 587 |
| 52 | Keyence_Nbenth_myc_64.jpg | 16632 x 5792 | 126 | 942 | 519 | 3 | 1241 |
| 53 | Keyence_Nbenth_myc_65.jpg | 16632 x 5792 | 126 | 561 | 426 | 12 | 781 |
| 54 | Keyence_Nbenth_myc_8.jpg | 9560 x 3204 | 126 | 269 | 243 | 14 | 202 |
| 55 | Keyence_Nbenth_myc_9.jpg | 9560 x 3205 | 126 | 254 | 246 | 11 | 204 |
| | | | **Count** | **20564** | **16420** | **508** | **25077** |

**Table S4. Test dataset.** M+: tiles labelled as colonised root sections; M−: tiles labelled as non-colonised root sections; Other: any other tile (background, debris, air bubbles, etc.); A: tiles containing arbuscules; V: tiles containing vesicles; H: tiles containing hyphopodia; IH: tiles containing intraradical hyphae; px: pixels; n.d.: not determined.

| # | Image name | Acquisition device | Image size (px) | Tile size (px) | M+ | M− | Other | A | V | H | IH |
|---|---|---|---|---|---|---|---|---|---|---|---|
| 1 | Large1_norm.jpg | Digital microscope | 1134x1134 | 126 | 21 | 5 | 55 | 16 | 15 | 0 | 18 |
| 2 | Large2_norm.jpg | Digital microscope | 1134x1134 | 126 | 44 | 6 | 31 | 28 | 31 | 2 | 36 |
| 3 | Large3_norm.jpg | Digital microscope | 1134x1134 | 126 | 19 | 8 | 54 | 10 | 12 | 0 | 17 |
| 4 | Large4_norm.jpg | Digital microscope | 1134x1134 | 126 | 38 | 3 | 40 | 28 | 31 | 1 | 33 |
| 5 | Large5_norm.jpg | Digital microscope | 1134x1134 | 126 | 30 | 4 | 47 | 26 | 22 | 0 | 26 |
| 6 | Large6_norm.jpg | Digital microscope | 1134x1134 | 126 | 40 | 9 | 32 | 37 | 30 | 2 | 23 |
| 7 | Large7_norm.jpg | Digital microscope | 1134x1134 | 126 | 33 | 6 | 42 | 31 | 24 | 1 | 18 |
| 8 | Large8_norm.jpg | Digital microscope | 1134x1134 | 126 | 26 | 20 | 35 | 21 | 18 | 1 | 25 |
| 9 | Large9_norm.jpg | Digital microscope | 1134x1134 | 126 | 20 | 13 | 48 | 12 | 10 | 0 | 19 |
| 10 | Large10_norm.jpg | Digital microscope | 1134x1134 | 126 | 14 | 20 | 47 | 6 | 4 | 0 | 14 |
| 11 | Small1_norm.jpg | Flatbed scanner | 360x360 | 40 | 32 | 15 | 34 | n.d. | n.d. | n.d. | n.d. |
| 12 | Small2_norm.jpg | Flatbed scanner | 360x360 | 40 | 30 | 9 | 42 | n.d. | n.d. | n.d. | n.d. |
| 13 | Small3_norm.jpg | Flatbed scanner | 360x360 | 40 | 37 | 18 | 26 | n.d. | n.d. | n.d. | n.d. |
| 14 | Small4_norm.jpg | Flatbed scanner | 360x360 | 40 | 34 | 10 | 37 | n.d. | n.d. | n.d. | n.d. |
| 15 | Small5_norm.jpg | Flatbed scanner | 360x360 | 40 | 22 | 17 | 42 | n.d. | n.d. | n.d. | n.d. |
| 16 | Small6_norm.jpg | Flatbed scanner | 360x360 | 40 | 15 | 37 | 29 | n.d. | n.d. | n.d. | n.d. |
| 17 | Small7_norm.jpg | Flatbed scanner | 360x360 | 40 | 36 | 2 | 43 | n.d. | n.d. | n.d. | n.d. |
| 18 | Small8_norm.jpg | Flatbed scanner | 360x360 | 40 | 42 | 13 | 26 | n.d. | n.d. | n.d. | n.d. |
| 19 | Small9_norm.jpg | Flatbed scanner | 360x360 | 40 | 18 | 15 | 48 | n.d. | n.d. | n.d. | n.d. |
| 20 | Small10_norm.jpg | Flatbed scanner | 360x360 | 40 | 27 | 18 | 36 | n.d. | n.d. | n.d. | n.d. |

**Table S5. CNN1 performance on various host plants.** M+: colonised root sections; M−: non-colonised root sections; Other: background and contaminants.

| Species | Accuracy (%) | | | Sensitivity (%) | | | Specificity (%) | | |
|---|---|---|---|---|---|---|---|---|---|
| | M+ | M– | Other | M+ | M– | Other | M+ | M– | Other |
| *M. truncatula* | 98 | 98 | 100 | 100 | 93 | 100 | 98 | 100 | 100 |
| *L. japonicus* | 96 | 92 | 96 | 90 | 77 | 97 | 98 | 96 | 93 |
| *O. sativa* | 97 | 96 | 99 | 95 | 75 | 98 | 99 | 97 | 100 |

**Table S6. CNN2 performance on various host plants.** A: arbuscules; V: vesicles; IH: intraradical hyphae.

| Species | Accuracy (%) | | | Sensitivity (%) | | | Specificity (%) | | |
|---|---|---|---|---|---|---|---|---|---|
| | A | V | IH | A | V | IH | A | V | IH |
| *M. truncatula* | 85 | 91 | 86 | 93 | 82 | 95 | 83 | 100 | 80 |
| *L. japonicus* | 84 | 88 | 95 | 98 | 95 | 98 | 80 | 73 | 92 |
| *O. sativa* | 86 | 92 | 91 | 90 | 85 | 89 | 83 | 92 | 94 |

**Table S7. CNN1 performance on various AM fungi.** M+: colonised root sections; M−: non-colonised root sections; Other: background and contaminants; n.d. : not determined (no background tiles).

| Species | Accuracy (%) | | | Sensitivity (%) | | | Specificity (%) | | |
|---|---|---|---|---|---|---|---|---|---|
| | M+ | M– | Other | M+ | M– | Other | M+ | M– | Other |
| *C. claroideum* | 95 | 92 | n.d. | 97 | 96 | n.d. | 90 | 96 | n.d. |
| *R. microaggregatum* | 100 | 92 | 92 | 100 | 100 | 79 | 100 | 91 | 100 |
| *F. geosporum* | 100 | 100 | n.d. | 100 | 100 | n.d. | 100 | 100 | n.d. |
| *F. mosseae* | 97 | 97 | 100 | 96 | 100 | 100 | 100 | 97 | 100 |

**Table S8. CNN2 performance on various AM fungi.** A: arbuscules; V: vesicles; IH: intraradical hyphae. n.d. : not determined.

| Species | Accuracy (%) | | | Sensitivity (%) | | | Specificity (%) | | |
|---|---|---|---|---|---|---|---|---|---|
| | A | V | IH | A | V | IH | A | V | IH |
| *C. claroideum* | 95 | 97 | 82 | 85 | 81 | 98 | 96 | 95 | 98 |
| *R. microaggregatum* | 92 | 84 | 74 | 94 | 97 | 90 | 80 | 84 | 72 |
| *F. geosporum* | 92 | 68 | 75 | 100 | 50 | 98 | 87 | 70 | 71 |
| *F. mosseae* | 93 | 70 | 79 | 98 | n.d. | 94 | 70 | 100 | 65 |
